# Differences in the genetic structure between and within two landlocked Ayu groups with different migration patterns in Lake Biwa revealed by environmental DNA analysis

**DOI:** 10.1101/2022.04.11.487953

**Authors:** Satsuki Tsuji, Naoki Shibata, Hayato Sawada, Katsutoshi Watanabe

## Abstract

Ayu, *Plecoglossus altivelis*, is largely an annual amphidromous fish, although a landlocked population lives in Lake Biwa, the largest lake in Japan. The landlocked population comprises two migrant groups, spring migrants and autumn migrants, which run to inlet rivers from the lake at different seasons. We used environmental DNA (eDNA) analysis, which is reported to be more sensitive and cost-effective than capture surveys, to clarify the genetic structure of this landlocked Ayu population with different migration patterns in Lake Biwa. We took water samples in 11 inlet rivers in the spring and autumn for two years in a row and quantitatively detected a total of 265 haplotypes of the mitochondrial D-loop region. The pairwise fixation index (*F*_ST_) value and haplotype diversity indicated that there were genetic differences between the two migrant groups in their respective rivers, and the *F*_ST_ values were negatively related to latitude and the presence of artificial fish release. Additionally, the isolation by distance within each migrant group was observed when the lake was divided into the east and west sides. These findings show that the landlocked Ayu population in Lake Biwa has the genetic structure associated with migration patterns and geographical distance. This study demonstrates that the eDNA approach will be effective for conducting a large-scale investigation of genetic structure beyond simple presence/absence tests.

## Introduction

Ayu (*Plecoglossus altivelis*; Plecoglossidae, Teleostei) is a keystone species in Japanese stream ecosystems, occupying a unique ecological niche as an algivorous specialist (Miyaji, 1960) and one of the most important commercial fishes in the Japanese freshwater fisheries (Iguchi et al., 2002; Kawanabe, 1996). Mostly, Ayu has an amphidromous life history, in which the larvae inhabit the coastal zone and migrate to rivers from the sea for further growth, maturation and reproduction throughout a one-year life cycle (Nishida, 2001). On the other hand, landlocked populations of this species are found in a few lakes, including Lake Biwa, the largest lake in Japan. The population in Lake Biwa is enormous, several hundred to 2,000 metric tonnes of this fish being annually yielded (Ministry of Agriculture, Forestry and Fisheries, 2021). Intriguingly, the majority of population’s life history is not amphidromous but anadromous-like (cf. Azuma, 1970, 1973a; Matsuyama and Matsuura, 1985); the majority of the fish enter inlet rives only for spawning.

The different life-history patterns in the landlocked Ayu population in Lake Biwa have long been of interest in the fields of fisheries, ecology, and evolution (Kawanabe, 1985, 1996). The population is typically divided into two migrant groups based on the run timing to inlet rivers from the lake (Fig. 1); (1) the spring migrants (late March–April; so-called “O-Ayu”), which grow in the middle to upper reaches of the inlets and spawn at the lower reaches (i.e. amphidromous-like) and (2) the autumn migrants (late August–September; so-called “Ko-Ayu”), which spend nearly their entire lives in the lake and migrate to the inlets to spawn in autumn (i.e. anadromous-like) (Azuma, 1973b; Iguchi et al., 2002; Iguchi and Nishida, 2000). With the exception that the nursery ground is a freshwater lake rather than the sea, the former’s life history is essentially identical to that of other river populations of this species. Spring migrants feed epilithic algae on the river bed (vs. zooplankton in the lake; Azuma, 1973a, 1973b) and finally grow much larger than the autumn migrants (standard length ± SD: spring migrants, 17.1 ± 0.7 cm; autumn migrants, 9.2 ± 0.4 cm; Iguchi 1996). On the other hand, the autumn migrants have much larger population size than the spring migrants (cf. Azuma, 1970, 1973a, 1973b, 1973c). Furthermore, the autumn migrants tend to spawn earlier (late August–late September) than the spring migrants (middle September–middle November) (Azuma, 1973b; Tsukamoto and Kajihara, 1987).

**Fig. 1.**
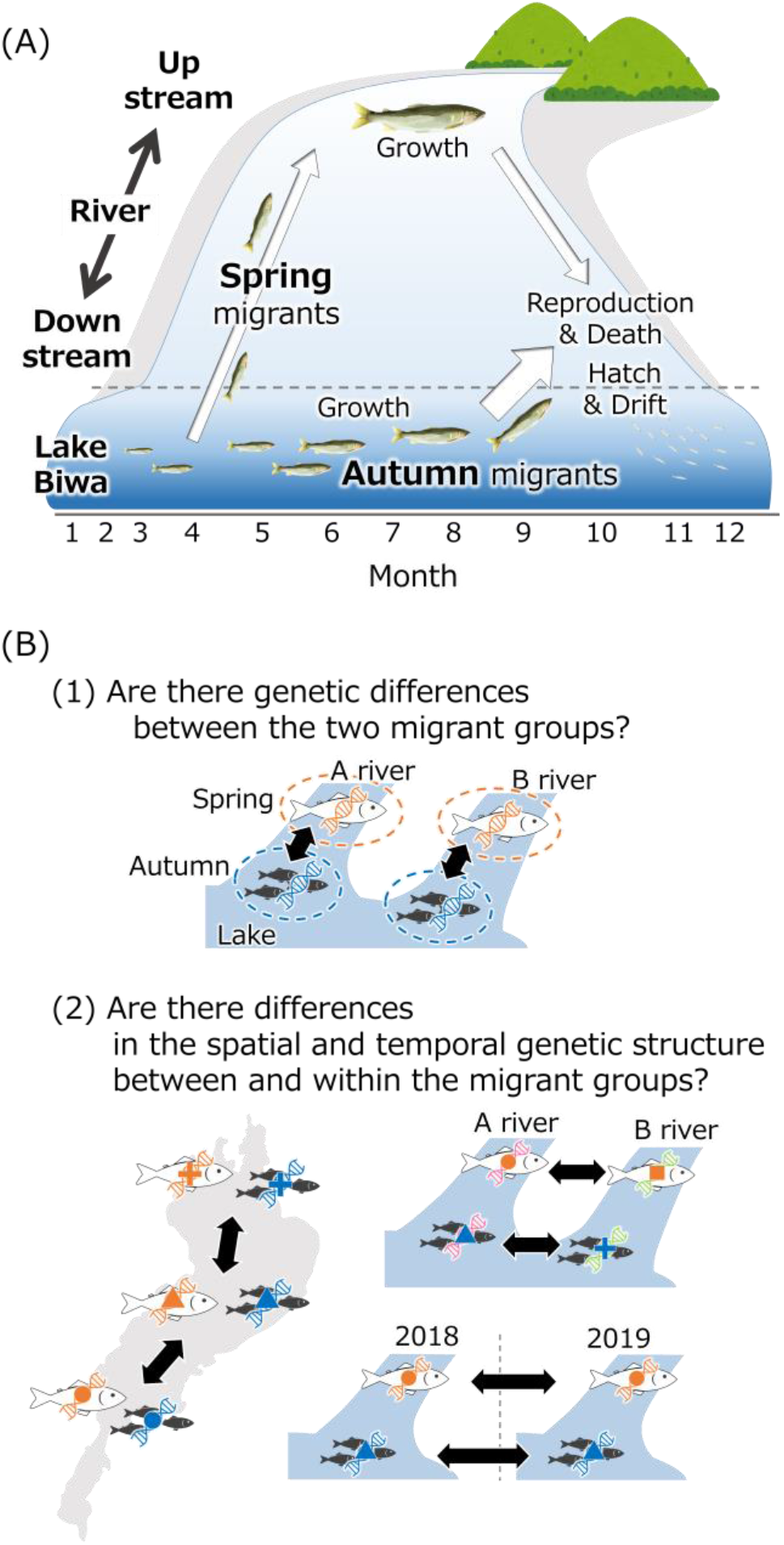
(A) Differences in the migration timing and growth areas in the two migrant groups (spring and autumn migrants) of the Ayu population landlocked in Lake Biwa and (B) the two hypotheses to be tested in this study.

Diversification of migration patterns often facilitates reproductive isolation between the migrant groups, which may lead to population differentiation (Moser et al., 2012). This could be due to several factors: (1) the difference in body size between different types could drive a sexual assortative mating through mate choice (Moser et al., 2012); (2) the shift in reproductive timing associated with the divergence could cause phenological assortative mating (e.g. Feder et al., 1997; Santos et al., 2011). These are both plausible in the case of Ayu in Lake Biwa; it tends to choose mating partners of the similar size as themselves (Iguchi and Maekawa, 1993; Nishida, 1974), and autumn migrants tend to spawn earlier than spring migrants (Azuma, 1973b; Tsukamoto et al., 1987). These facts lead to the idea that some extent of reproductive isolation has developed between the two migrant groups and they might be in the early phases of population differentiation. If it is the case, the landlocked Ayu in Lake Biwa will provide an excellent opportunity to observe the process of population differentiation that occurs as a result of landlocking.

The only prior study that investigated the genetic variation between and within the two migrant groups, found that there were tiny but significant genetic differences between the two groups (*F*_ST_ < 0.34 in the mtDNA D-loop region; Iguchi et al. 2002). Also, within the autumn migrants, there was tendency of isolation by distance. However, because of the limited sample size in both the specimens and rivers analysed, further study with more sophisticated sampling designs both spatially and temporally is needed (Iguchi et al. 2002). If the landlocked Ayu in Lake Biwa is not genetically homogeneous as suggested by Iguchi et al. (2002), it should be considered in fisheries management including fishing, stocking and seed production. For example, although the fish stocking is an effective way to reduce anthropogenic impacts on fishery critical resources, it could also impose ecological and genetic risks if genetically improper fish are stocked (Iguchi, 2011; Laikre et al., 2010). In Lake Biwa, a large number of the juveniles captured from the lake are stocked in the middle to upper reaches of several inlet rivers every year in early May without consideration of the genetic aspects (Azuma, 1973b; Iguchi et al., 2002). Currently, the magnitude of impacts of stocking on population dynamics and genetic structure of the population cannot be accurately examined due to a lack of information about the exact genetic structure of landlocked Ayu population in Lake Biwa.

The purpose of this study was to clarify the genetic structure of the landlocked Ayu population in Lake Biwa with different migration patterns. To perform a rapid and geographically wide-ranging survey, we adopted environmental DNA analysis (eDNA) for the evaluation of genetic diversity and differentiation. The eDNA analysis is employed as a highly-sensitive and cost-effective tool for species detection (Rees et al., 2014). Furthermore, it has recently started to be used to evaluate intraspecific genetic diversity within and among populations (Sigsgaard et al., 2020; Tsuji et al., 2020a). For example, an eDNA-based mtDNA survey using one-litre water samples provided comparable results from the conventional capture-based method with the Sanger sequencing for a large number (∼100) of specimens (Tsuji et al., 2020b). In addition, an eDNA-based survey enables reasonable estimation of genetic indicators (e.g. nucleotide diversity, Tajima’s test statistic, and effective female population size) by using the relative number of DNA copies of each mtDNA haplotype amplified from water samples (Tsuji et al., 2020c). To achieve the goal of this study, we test the following two hypotheses using population genetic parameters estimated by eDNA analysis: (1) whether there are genetic differences between the two migrant groups, and (2) whether there are differences in the spatial and temporal genetic structure between the migrant groups. Based on results, we explored the factors of genetic differentiation between migrant groups and geographical population structure, with particular attention to spawning timing and lake currents.

## Materials and Methods

### Study sites

The genetic structure of the two migration groups of the landlocked Ayu was examined using an eDNA-based method in 11 inlet rivers of Lake Biwa in Shiga Prefecture, central Honshu, Japan. The Echi River (1-EC), the Uso River (2-US), the Inukami River (3-IN), the Seri River (4-SR), the Amano River (5-AM), the Shiotsu-o River (6-SO), the Chinai River (7-CN), the Ishida River (8-IS), the Ado River (9-AD), the Wadauchi River (10-WD) and the Wani River (11-WN) were all sampled in the lower reaches of each river (Fig. 2). A large number of juveniles captured from Lake Biwa were stocked into the middle and/or upper reaches in 1-EC, 3-IN, 8-IS and 9-AD every spring. Until 2008, stocking was done in 4-SR.

**Fig. 2.**
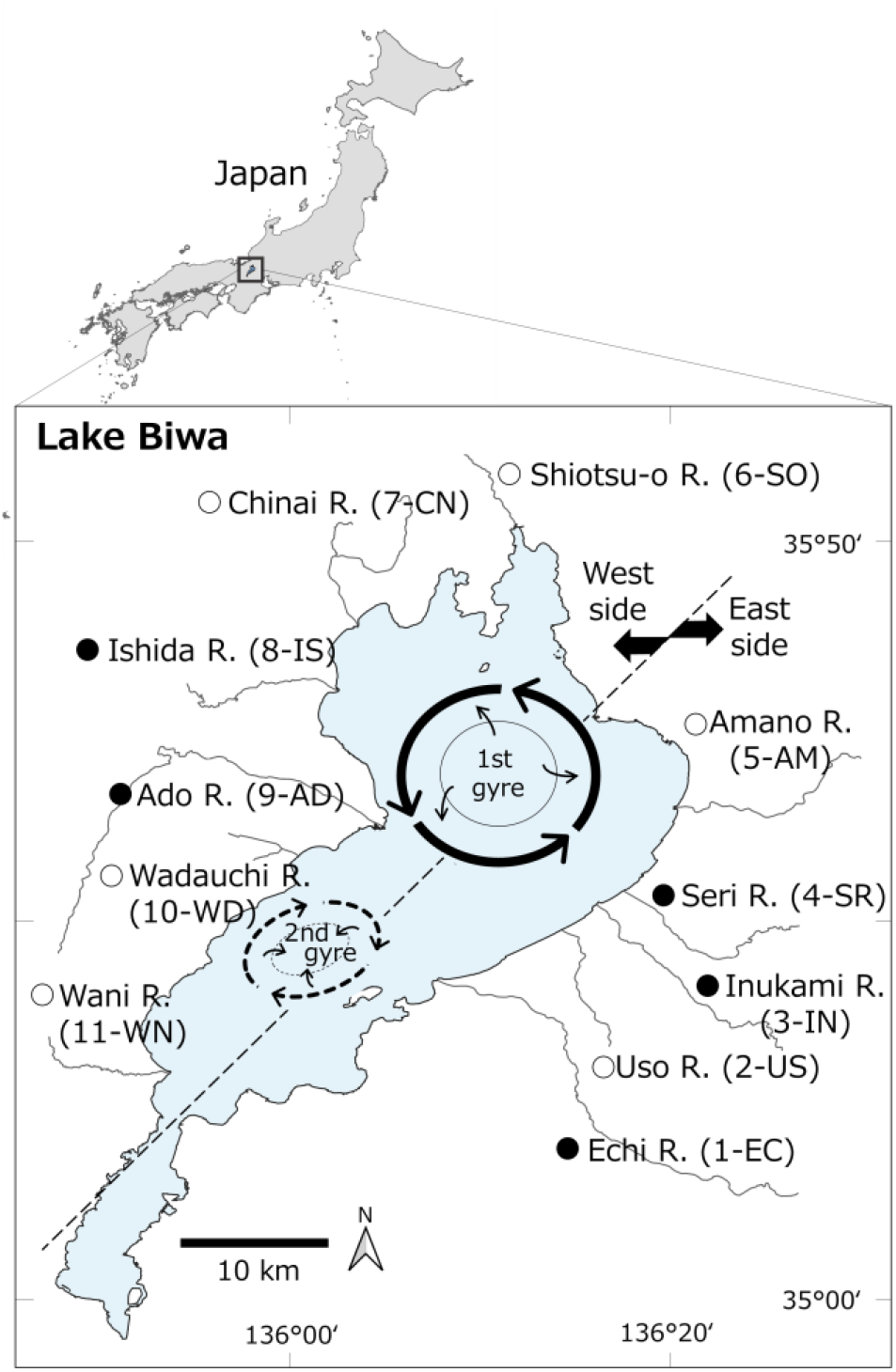
The studied rivers and gyre system in Lake Biwa. The solid and open circles indicate the rivers where fish have been stocked or not, respectively.

### Water sample processing and eDNA extraction

Water sampling was performed in each river in spring and autumn in 2018 and 2019 (April 22 and September 14, 2018 and April 19 and August 27, 2019). By collecting water samples from the lower reaches of rivers at the time when each migrant group migrates (i.e. Spring or Autumn), we can collect eDNA derived from individuals belonging to each group. Autumn water sampling was performed before the spring migrants moved from the upper to lower reaches of the river to spawn. A total of 4 L of surface water was collected from three to five points in the transverse direction at each sampling site using a disposable plastic cup. The collected water was pooled and carefully agitated with a bucket. The water qualities of each sample are listed in Table S1. The 3 L of water sample was filtered using three Whatman GF/F filters (1 L/filter; GE Healthcare, Japan) on-site. To monitor contamination during the filtering and subsequent DNA extraction steps, 1 L of ultrapure water was filtered on-site in the same manner. The filter samples were placed in a plastic bag and immediately kept at -20 °C. All filtering equipment was decontaminated by soaking in the 10% bleach solution water before use. DNA was extracted from each filter sample using the spin column (EP-31201, GeneDesign, Japan) and DNeasy Blood and Tissue Kit (Qiagen, Germany). At the final extraction step, DNA was eluted from the column with 100 µL of Buffer AE. Detailed information about DNA extraction is described in Appendix.

### Paired-end library preparation and sequencing with MiSeq

To quantitatively evaluate the concentration of each Ayu haplotype, we adopted the quantitative eDNA metabarcoding method using the internal standard DNAs (qMiSeq method, Ushio et al., 2018). The use of the qMiSeq method enables us to estimate eDNA copy numbers of detected unknown Ayu haplotypes by using sample-specific standard lines which were obtained for each sample based on the relationship between the number of sequence reads and the known copy number of standard DNAs (Table S2: Tsuji et al., 2020c; Ushio et al., 2018). In the DNA library preparation, we employed a two-step tailed PCR approach to construct the paired-end libraries.

The first-round PCR (1st PCR) was carried out with a 12-μl reaction volume to amplify the Ayu mitochondrial D-loop region and the standard DNAs (insert length, 166 bp, both). The sequences of primers are: PaaDlp-2_F (5′-ACA CTC TTT CCC TAC ACG ACG CTC TTC CGA TCT NNN NNN CCG GTT GCA TAT ATG GAC CTA TTA C-3′), PaaDlp-2_R1 and R2 (5′- GTG ACT GGA GTT CAG ACG TGT GCT CTT CCG ATC TNN NNN NGC TAT TRT AGT CTG GTA ACG CAA G -3′). The second-round PCR (2nd PCR) was carried out with a 12-μl reaction volume to add Illumina sequencing adaptors and the dual-index (i.e. barcode) sequences. The sequences of primers are: Forward (5′-AAT GAT ACG GCG ACC ACC GAG ATC TAC AXX XXX XXX ACA CTC TTT CCC TAC ACG ACG CTC TTC CGATCT-3′), and Reverse (5′-CAA GCA GAA GAC GGC ATA CGA GAT XXX XXX XXG TGA CTG GAG TTC AGA CGT GTG CTC TTC CGA TCT-3′).

All indexed products of the second PCR were pooled in equal volumes, and the target size of the libraries (ca. 370-bp) was excised using 2% E-Gel SizeSelect Agarose Gels (Thermo Fisher Scientific, USA). The excised library was adjusted to 4 nM and sequenced on the MiSeq platform using MiSeq v2 Reagent Micro or Nano Kit for 2 × 150 bp PE cartridges (Illumina, San Diego, USA) with 30% PhiX spike-in according to the manufacturer’s instructions. All raw sequences obtained in this study were deposited in the DDBJ Sequence Read Archive (accession number: DRA013835). Detailed information about paired-end library preparation and sequencing by MiSeq are described in Appendix.

### Bioinformatic analysis

To perform the denoising of erroneous sequences, the Fastq files containing raw reads were processed using the Divisive Amplicon Denoising Algorithm 2 package ver. 1.12.1 (DADA2, Callahan et al., 2016) of R by each MiSeq run. The denoising was performed by a method used in a previous studies (Tsuji et al., 2020a, 2020b, 2020c). Briefly, first, the “dada2::filterAndTrim” function was used for the removal of excess sequences (primers and random hexamers), and the trimming and filtering of the forward and reverse reads. The error model was trained using the ‘dada2::learnErrors’ function and used to identify and correct indel-mutations and substitutions of dereplicated passed sequences using the “dada2::dada” function. The paired reads were merged, and then the sequences were assigned to each standard DNA sequence or each unknown haplotype using the function “dada2::assignSpecies”. The parameters that control the stringency of the denoising step in DADA2, OMEGA_A and OMEGA_C, were set at the default values. To convert the number of sequence reads to the number of DNA copies, a linear regression line between sequence reads of standard DNAs and their known DNA copies was obtained as a standard line for each sample using the ‘lm’ function in R (the intercept was set to 0). For each sample, the number of DNA copies of detected unknown haplotypes was calculated as *Nc = MRs/S* using the obtained standard line (*MRs* = the number of MiSeq sequence reads, *S* = the regression slope of the sample-specific standard line). The calculated *Nc* values are hereafter referred to as “calculated copy numbers”. The detailed workflow of qMiSeq is described in the Ushio et al. (2018) and Tsuji et al. (2020c).

### Data screening

To remove chimeric sequences derived from the standard DNAs, the detected haplotypes containing specific sequences to the standard DNAs (std1, 5′-ACA TAT AGT AGT GAG AAC CCA CCA ACT GAT GAT-3′; std2, 5′-GGG ATA ACC ATC CCC TAT ATG GTT TAG TAC ATA-3′; std3, 5′-TGA GAG ACC ACC AAC TGA TTT GTA TAA AGG TAC-3′; Tsuji et al. 2020c) were manually searched and removed. Additionally, the accuracy of eDNA-based detection of haplotypes would be further increased by selecting only haplotypes detected from all filtration replicates because the error sequences are generated randomly and accidentally in each sample (Tsuji et al. 2020b). Thus, only the haplotypes detected in all three filtration replicates and the average number of sequence reads of them were used for subsequent analysis. To further improve the detection accuracy, the haplotypes detected with an average of less than one copy per litre were removed. The core algorithm of DADA2 merges the sequence reads inferred as errors with the most similar sequence to each (see Callahan et al. 2016 for the detailed algorithm of DADA2). Thus, the sequences with many similar sequences that were recognized as errors may be overestimated in the number of reads. To reduce the observation error, if the coefficient of variation of the observed number of sequence reads among three filtration replicates was exceeded one, an outlier was removed and the average value in the other two replicates was used for subsequent analysis. Previous studies have employed this data screening method and obtained results comparable to large-scale capture surveys (96 individuals/site) (Tsuji et al., 2020b).

In addition, considering the high genetic diversity of the landlocked Ayu population in Lake Biwa, at least multiple haplotypes are expected to be detected in each sample (Iguchi et al., 2002; Tsuji et al., 2020b and 2020c). If the number of detected haplotypes was below that of the minimum number in previous study (5 haplotypes; Iguchi et al. 2002), the sample was excluded from subsequent analyses because they were suspected of having false negative haplotypes due to low eDNA concentrations. Furthermore, in the autumn survey in 2018, the water level in all rivers was still higher than the normal water levels due to the typhoon that passed through a few days before the survey. The range of water level rise of each river was as follows: 1–5 cm, 2-US, 5-AM and 7-CN; 6–10 cm, 6-SO; 11–15 cm, 3-IN and 8-IS; 16–20 cm, 4-SR and 11-WN; >20, 1-EC and 9-AD; and not available for 10-WD (data provided from Flood Management Office of Shiga Prefecture). Especially, in 1-EC and 9-AD, where the water level was particularly high, the number of detected haplotypes were obviously lower than in 2018 spring (see Results, Table 1), even though the autumn migrants have significantly larger biomass than that of spring migrants (cf. Azuma, 1970, 1973a, 1973b and 1973c). Thus, the decrease in the number of detected haplotypes was considered to be more strongly influenced by the dilution due to rising water levels than other rivers. For that reason, 1-EC 2018-autumn and 9-AD 2018-autumn were excluded from subsequent analyses.

**Table 1.** Number of detected haplotypes and estimated haplotype diversity (*h*) for each sample. The data with one or two asterisks were excluded from analysis because of the presence of suspected false negative haplotypes due to the low eDNA concentration. NA indicates cases where the water sampling could not be performed due to river construction.

| River | Basin area<br>(km<sup>2</sup>) | Number of detected haplotypes | | | | Haplotype diversity ($h$) | | | |
| --- | --- | --- | --- | --- | --- | --- | --- | --- | --- |
|  |  | 2018 |  | 2019 |  | 2018 |  | 2019 |  |
|  |  | Spring | Autumn | Spring | Autumn | Spring | Autumn | Spring | Autumn |
| 1-EC | 204.0 | 22 | 13** | 26 | 47 | 0.83 | NA | 0.87 | 0.88 |
| 2-US | 69.7 | 1* | 40 | 7 | 62 | NA | 0.87 | 0.78 | 0.89 |
| 3-IN | 101.0 | 43 | 131 | 16 | 81 | 0.89 | 0.90 | 0.84 | 0.90 |
| 4-SR | 65.0 | 46 | 141 | 14 | 95 | 0.89 | 0.91 | 0.75 | 0.92 |
| 5-AM | 109.0 | 9 | 89 | 1* | 57 | 0.82 | 0.89 | NA | 0.90 |
| 6-SO | 19.6 | 39 | 135 | 38 | 56 | 0.88 | 0.91 | 0.91 | 0.88 |
| 7-CN | 51.0 | 84 | 136 | 71 | 85 | 0.89 | 0.91 | 0.92 | 0.90 |
| 8-IS | 59.0 | 53 | 156 | NA | NA | 0.89 | 0.91 | NA | NA |
| 9-AD | 306.0 | 91 | 8** | 53 | 68 | 0.89 | NA | 0.77 | 0.89 |
| 10-WD | 10.0 | 8 | 31 | 9 | 50 | 0.69 | 0.86 | 0.77 | 0.89 |
| 11-WN | 16.0 | 9 | 80 | 1* | NA | 0.72 | 0.89 | NA | NA |
| Average |  |  |  |  |  | 0.84 | 0.89 | 0.83 | 0.89 |
| SD |  |  |  |  |  | 0.07 | 0.02 | 0.07 | 0.01 |

### Population genetic analysis

As an indicator to represent intraspecific diversity, haplotype diversity (*h*; Nei and Tajima 1981) was estimated according to the following formula: 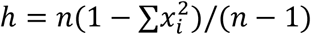, where *n* is the calculated copy number of each detected haplotype, *x*_*i*_ is the frequency of the calculated copy number of haplotype *i* relative to the total copy number. Furthermore, the pairwise fixation index values (*F*_ST_) were estimated using Arlequin ver. 3.5.2.2 (Excoffier and Lischer, 2010) based on the calculated copy number of each detected haplotype to compare the genetic structure among different sample groups (i.e., between migrant groups for each river and between geographic samples of each migrant group).

### Statistical analysis

To confirm the validity of the estimated haplotype diversity using eDNA analysis, the mean values of the estimated *h* of autumn migrants was compared with that of a previous study based on a capture survey (average *h* 0.78, standard deviation 0.19; Iguchi et al. 2002) using the Kruskal-Wallis test (kruskal.test function in R). The difference in the number of detected haplotypes and the estimated haplotype diversity was tested between and within the migrant groups using the Wilcoxon signed-rank test (exactRankTests package ver. 0.8-32). In addition, since the basin area of each river may affect the carrying capacity of the Ayu population, the influence of the basin area on the number of detected haplotypes was tested using a generalized linear mixed-effects model (GLMM; glmmML package ver. 1.1.1) with a Poisson error distribution. In the GLMM model, the study rivers were treated as random effects. Also, a beta regression model (betareg package ver. 3.1-4) was used to investigate the effects of the latitude, longitude and presence/absence of fish stock on *F*_ST_ between two migrant groups in each river.

To investigate the spatial genetic structure within each migrant group, the relationship between *F*_ST_ values and the geographical distance for all sample pairs in each migrant group was assessed using the Mantel correlation test (ecodist package ver. 2.0.7). As a geographical distance among river mouths, three measurement methods were used: (1) the shortest straight lines distance through the lake, (2) right-handed distance along the lakeshore, and (3) the shortest distance along the lakeshore. Additionally, assuming that Ayu does not migrate across the lake, we also examined the relationship between *F*_ST_ values and the shortest straight lines distance through the lake between rivers with dividing Lake Biwa into east and west (Fig. 2). Furthermore, the principal coordinate analysis (PCoA) based on *F*_ST_ matrix was performed to investigate the genetic structure between two migrant groups and within each migration group (cmdscale function in R). To examine whether the same river samples in different years were plotted closer to each other than those of different rivers, *F*_ST_ values were compared among the following four groups of pairs using the Kruskal-Wallis test (kruskal.test function in R) followed by Conover’s tests with Holm adjustment (PMCMRplus package ver.1.9.3); (1) same river pairs between 2018 and 2019, (2) different river pairs between 2018 and 2019, (3) different river pairs of 2018, and (3) different rivers pairs of 2019. If the samples came from the same rivers in different years are genetically close, their *F*_ST_ value of group (1) is expected to be smaller than those of the other groups. R version 4.1.1 software (R Core Team. R, 2019) was used for all statistical analyses, and the minimum level of significance was set at α = 0.05.

## Results

### MiSeq sequencing and data screening

The qMiSeq analysis with paired-end sequencing of total 153 libraries (132 real samples, 12 filter negative controls and 9 PCR negative controls) was successfully performed (total five MiSeq runs, Q30 > 96%). *R*^2^ values of the sample-specific standard line ranged from 0.88 to 1.00, and the median was 0.94 (Table S3). A total of 695 haplotypes were detected throughout the two-years survey at 11 rivers. Nucleotide substitution occurred at 63 sites out of 166 bp sequences. The number of sequence reads of detected haplotypes was converted to the copy number using the coefficient value of the sample-specific regression line (Table S3). After screening data, 265 haplotypes which were detected from all filtration replicates in each river in each season with one or more copies per litre were finally selected (Table S4).

As a general trend, the number of detected haplotypes for each river ranged from one to 156 (Table 1), with the majority of haplotypes shared among samples from different rivers and/or migrant groups. The data of 2-US (2018-spring), 5-AM (2019-spring) and 11-WN (2019-spring) in which only one haplotype was detected were excluded from subsequent analyses because they suspectedly have many false negative haplotypes (Table 1). Additionally, 1-EC (2018-autumn) and 9-AD (2018-autumn) were also excluded from subsequent analyses because they were suspected to have many false negative haplotypes due to the rise of the rivers (see Materials and Methods, Table 1).

### The number of detected haplotypes and haplotype diversity

The number of detected haplotypes significantly differed between spring and autumn migrants (Wilcoxon signed-rank test, *p* < 0.01 in both years), and more haplotypes were detected from the autumn migrants than the spring migrants in both years. On the other hand, there was no significant difference in the number of detected haplotypes between the different year samples of the same migrant group (Wilcoxon signed-rank test, *p* = 0.125 for spring, *p* = 0.078 for autumn). The GLMM results showed that the number of detected haplotypes was significantly increased by basin area (*p* < 0.001).

Haplotype diversity (*h*) of each sample ranged from 0.69 in 10-WD (2018-spring) to 0.92 in 7-CN (2019-spring) for the spring samples and 0.86 in 10-WD (2018-autumn) to 0.92 in 4-SR (2019-autumn) for the autumn samples (Table1). There was no significant difference between the mean estimated haplotype diversity of autumn migrants based on eDNA analysis and that in previous studies based on capture survey by Iguchi et al. (2002) (Kruskal-Wallis test, *p* = 0.75). The *h* values significantly differed between the spring and autumn migrants only in 2018 (Wilcoxon signed-rank test, *p* < 0.01 for 2018, *p* = 0.078 for 2019), but the autumn migrants showed higher *h* than that of spring migrants in almost all rivers in both years (8/8 rivers for 2018, 6/8 rivers for 2019, except for NA). On the other hand, there was no significant difference in *h* between the different year samples of the same migrant group (Wilcoxon signed-rank test, *p* = 0.66 for spring, *p* = 0.66 for autumn).

### Spatio-temporal patterns of genetic differences between the two migrant groups

In each river, the *F*_ST_ values between two migrant groups in each river varied from 0.0014 (7-CN 2018) to 0.030 (10-WD 2019) (Fig. 3 and Table S5). All of the calculated *F*_ST_ values were low, yet they differed significantly from zero (*p* < 0.001). The beta regression model showed a significant negative effect of latitude and the presence of fish stocking on the *F*_ST_ values between migrant groups in each river (coefficient estimates, -3.16 and -1.04; *p* < 0.05 and *p* < 0.01, respectively).

**Fig. 3.**
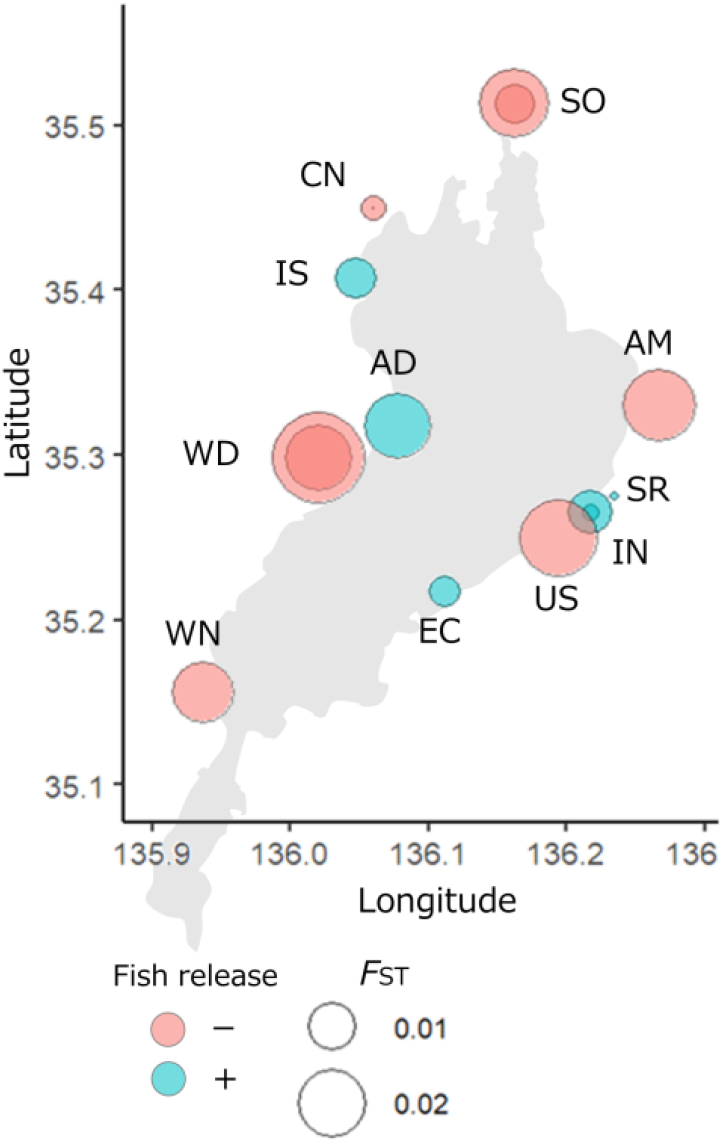
The relationship between the geographical coordinates of the sampling sites and the genetic distance (*F*_ST_) between two migrant groups in each river. The size and colour of the circles indicate the magnitude of *F*_ST_ and the presence (+) or absence (–) of fish stoking, respectively. *F*_ST_ was estimated by comparing different migrant groups in the same year for each river. The data for two years (if available) were shown as overlapping circles.

In both years, there was no significant association between *F*_ST_ within each migrant group and three measurements of geographical distance between the rivers when all rivers are taken into account (Mantel test, *p* > 0.05; Table 2 and Fig. 4). On the other hand, when Lake Biwa was divided into east and west sides and analysed separately, a significant association between *F*_ST_ value and the shortest straight lines distance was found in the rivers on the west side for spring migrants in 2018 (*p* < 0.05; Table 2 and Fig. 4). Although not statistically significant, the east side on both groups in 2018 and the west side on both groups in 2019 also showed high Mantel statistic r (0.543– 0.943).

**Table 2.**
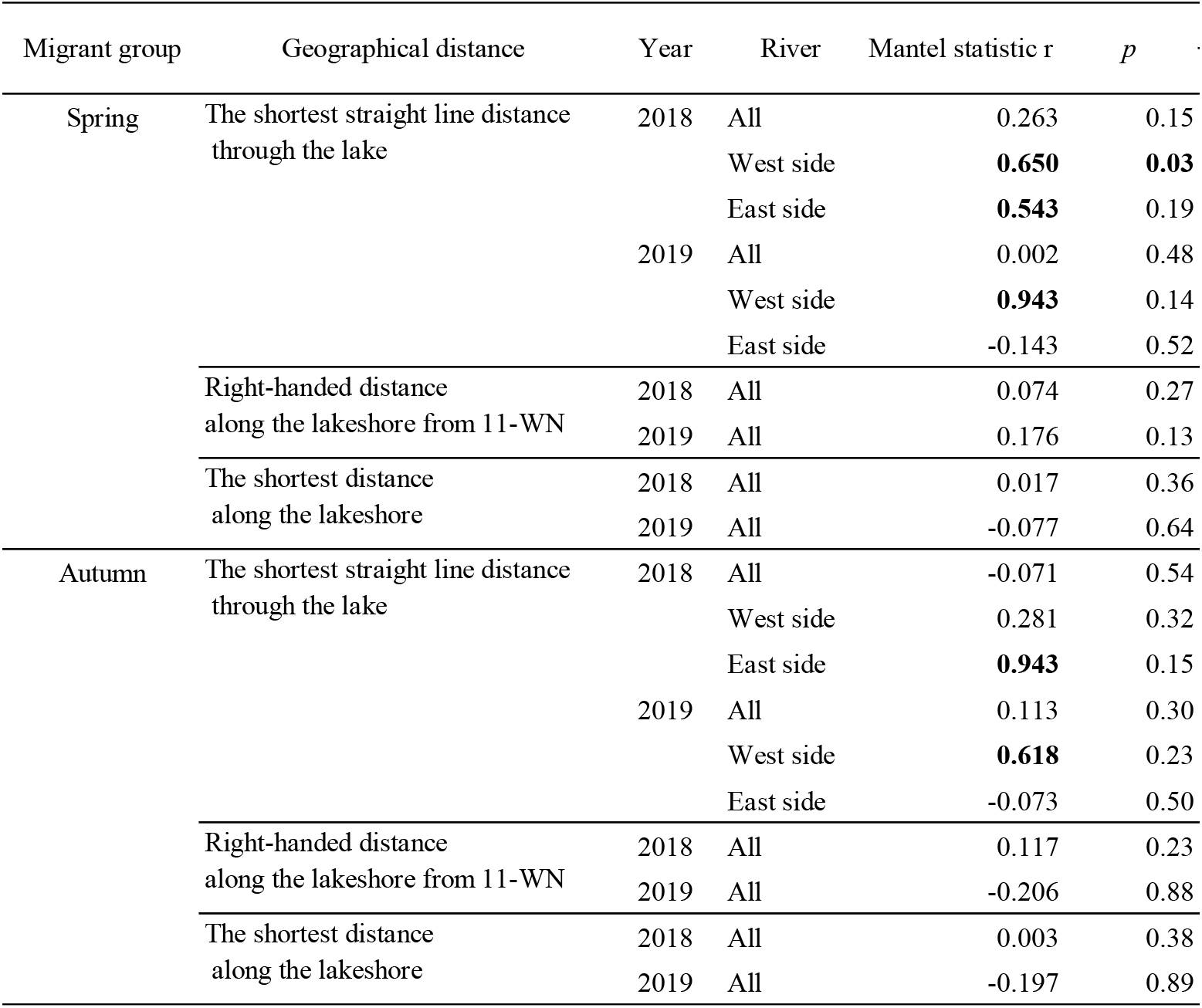
Results of Mantel tests for associations between the genetic distance (*F*_ST_) and the three measurements of geographical distance between study rivers. Bold letters indicate the statistical significances (*p* < 0.05) and the higher Mantel statistic r (*r* > 0.5)

| Migrant group | Geographical distance | Year | River | Mantel statistic $r$ | $p$ |
| --- | --- | --- | --- | --- | --- |
| Spring | The shortest straight line distance through the lake | 2018 | All | 0.263 | 0.15 |
|  |  |  | West side | **0.650** | **0.03** |
|  |  |  | East side | **0.543** | 0.19 |
|  |  | 2019 | All | 0.002 | 0.48 |
|  |  |  | West side | **0.943** | 0.14 |
|  |  |  | East side | -0.143 | 0.52 |
|  | Right-handed distance along the lakeshore from 11-WN | 2018 | All | 0.074 | 0.27 |
|  |  | 2019 | All | 0.176 | 0.13 |
|  | The shortest distance along the lakeshore | 2018 | All | 0.017 | 0.36 |
|  |  | 2019 | All | -0.077 | 0.64 |
| Autumn | The shortest straight line distance through the lake | 2018 | All | -0.071 | 0.54 |
|  |  |  | West side | 0.281 | 0.32 |
|  |  |  | East side | **0.943** | 0.15 |
|  |  | 2019 | All | 0.113 | 0.30 |
|  |  |  | West side | **0.618** | 0.23 |
|  |  |  | East side | -0.073 | 0.50 |
|  | Right-handed distance along the lakeshore from 11-WN | 2018 | All | 0.117 | 0.23 |
|  |  | 2019 | All | -0.206 | 0.88 |
|  | The shortest distance along the lakeshore | 2018 | All | 0.003 | 0.38 |
|  |  | 2019 | All | -0.197 | 0.89 |

**Fig. 4.**
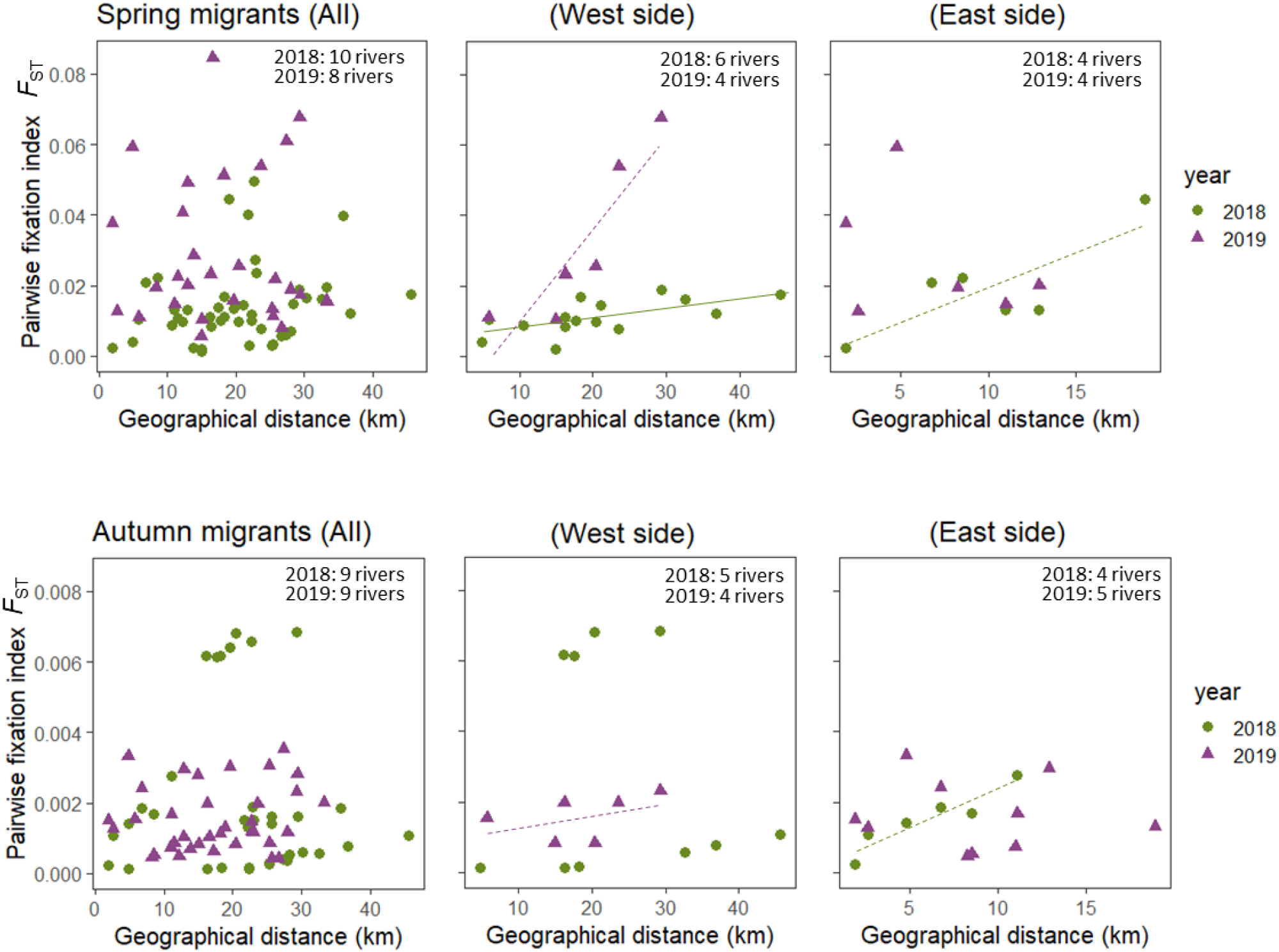
Relationship between the geographical distance (the shortest straight-line distance through the lake) and the genetic distance (*F*_ST_) among river samples for each migrant group. The green circle and purple triangle indicate data for 2018 and 2019, respectively. All, west side and east side indicate the results for data from all rivers studied, rivers on the western side and rivers on the eastern side of Lake Biwa, respectively. The solid and dashed lines indicate statistically significant relationships and the relationships that were not statistically significant but have a higher Mantel statistic *r* value (*r* > 0.5), respectively.

When the *F*_ST_ values of all sample pairs for each year were ordinated using a PCoA, spring migrants samples were spread widely, whereas autumn migrants samples were clustered in almost the same position, except for 10-WD (2018) (Fig. 5a, b). On the other hand, when the *F*_ST_ values among two-year samples for each migrant group were ordinated using a PCoA, plots of each river did not tend to cluster closely together (Fig. 5c, d). The *F*_ST_ comparisons also supported this result; *F*_ST_ values between the different year samples from same rivers were not smaller but similar to or rather larger than those of the other groups of pairs (Kruskal-Wallis test followed by Conover’s tests, *p* ≥ 0.05; Fig. 6). The comparisons of the *F*_ST_ values showed the significant differences between some groups in spring migrants (*p* < 0.01 for different rivers of 2018 vs. different rivers of 2019, *p* <0.001 for different rivers of 2018 vs. different rivers of 2018 and 2019; Fig. 6).

**Fig. 5.**
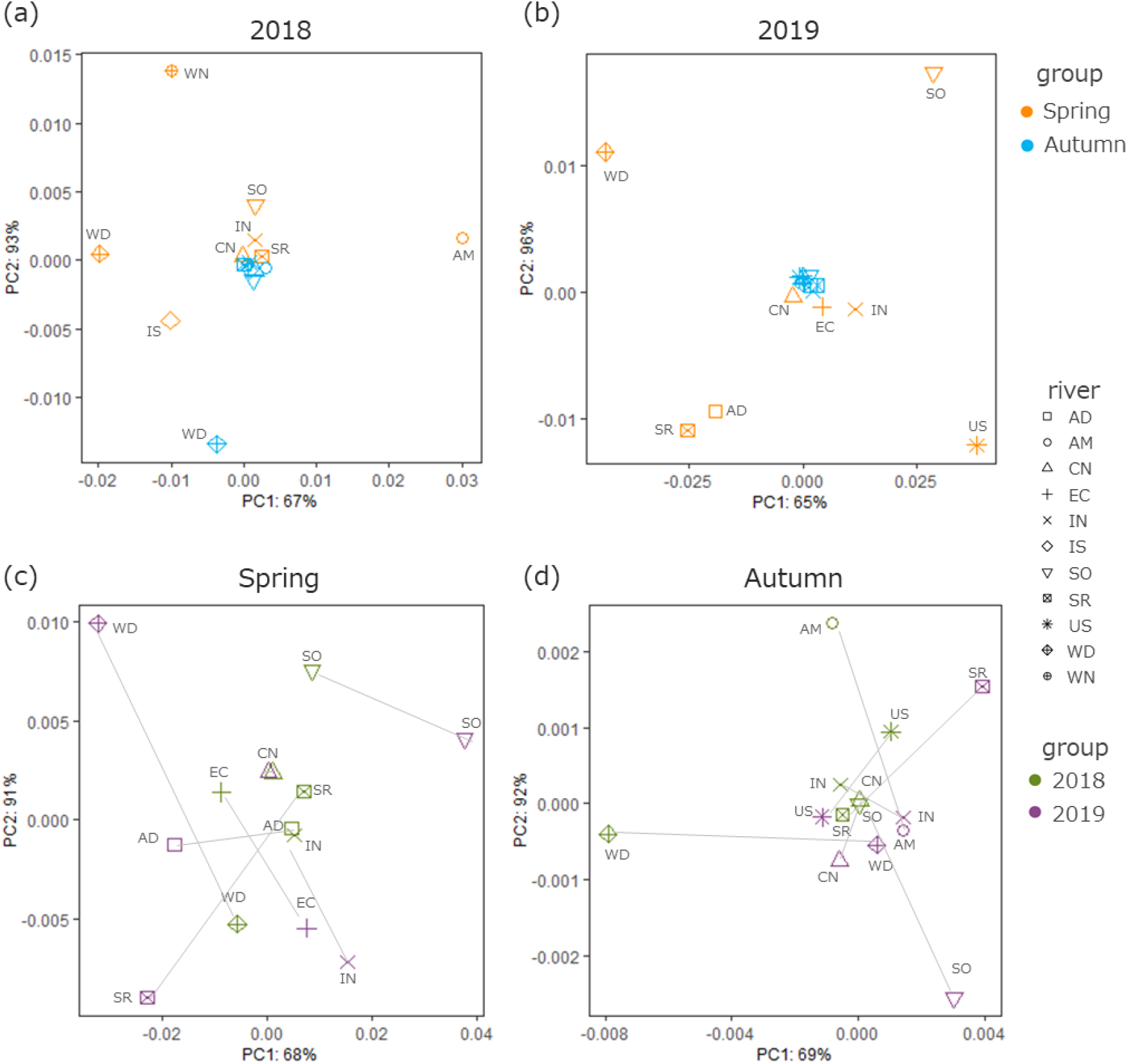
Results of the principal coordinates analysis (PCoA) on the genetic distance (*F*_ST_) between samples for each year (2018 and 2019 panels) and between samples for each migrant group (spring and autumn panels). The shapes of each plot indicate each river and are common to all panels. In the 2018 and 2019 panels, orange and blue indicate the data of spring migrants and autumn migrants, respectively. In spring and autumn panels, green and purple indicate data for 2018 and 2019, respectively. In spring and autumn panels, the plots of the same rivers are connected by lines.

**Fig. 6.**
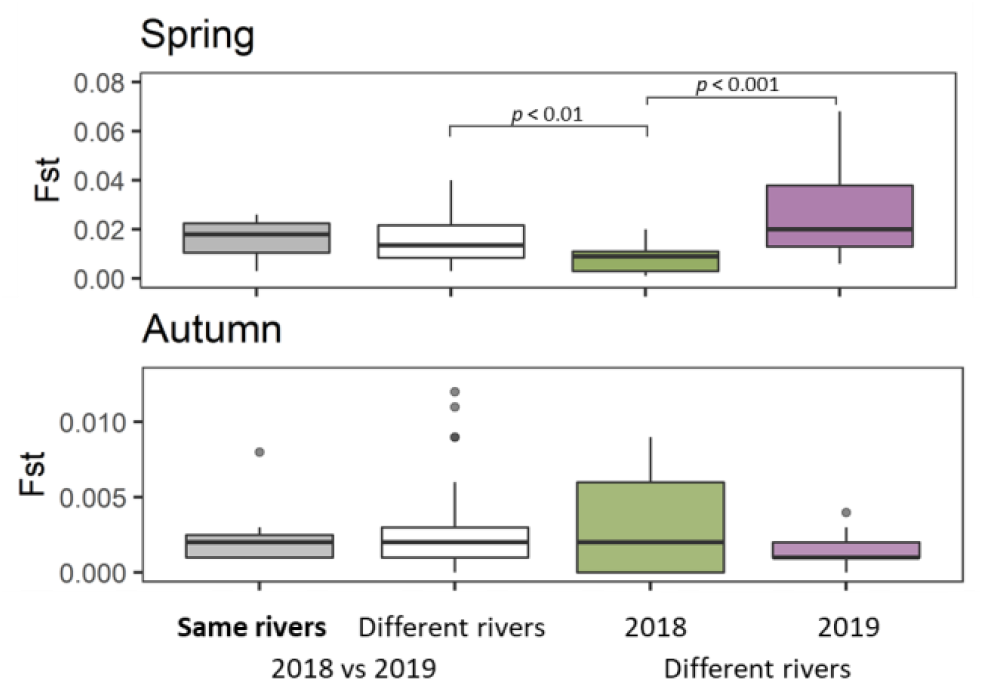
Comparisons of *F*_ST_ values among the following four groups of pairs; same rivers of 2018 and 2019, different rivers of 2018 and 2019, different rivers of 2018, and different rivers of 2019.

## Discussion

### The effectiveness and usefulness of eDNA analysis in the assessment of genetic diversity

Through the two sampling years, we successfully detected and quantified a large number of Ayu haplotypes (total 265) from 11 rivers by using eDNA analysis. Iguchi et al. (2002), the only previous study that investigated genetic variation between and within the spring and autumn migrant groups, analysed 15 specimens from each of five rivers, two of which were used for comparisons between the migrant groups. The study detected 38 haplotypes in a total of 120 specimens, which is significantly less than our findings. The application of eDNA analysis in our study allowed us to conduct a larger-scale and more extensive inquiry, which Iguchi et al. (2002) had identified as a potential problem. Furthermore, using the accurate estimated standard line for each sample, we quantitatively examined haplotype frequency and genetic diversity and successfully obtained reasonable *h* values that were consistent with the previous study based on capture survey. As also suggested by Tsuji et al. (2020c), the quantitative assessment of genetic diversity based on eDNA analysis enables the application of traditional population genetics tools/indicators. Because the statistical significance of those indicators is dependent on the sample size (in this case, the number of copies per litre), this point should be noticed and is examined in the next section.

### The genetic differences between the two migrant groups

Our *F*_ST_ assessment using eDNA revealed genetic differences between spring and autumn migrant groups in respective rivers, although the estimated *F*_ST_ values were generally small (0.001–0.019 for 2018, 0.003–0.030 for 2019). Additionally, autumn migrants possessed a larger number of haplotypes and showed higher haplotype diversity than spring migrants. This seems reasonable because the former have a much larger abundance than the latter (cf. Azuma, 1970,1973a, 1973b and 1973c) and supports that spring and autumn migrants are derived from groups with some degree of genetic differences. The small but significant genetic differences between the two migrant groups recognised in this study are consistent with the findings by the capture-based method of Iguchi et al. (2002), which reported 0.063–0.342 of *F*_ST_ for two river samples. The degree of freedom was not determinable for the population statistics calculated from eDNA data. However, at least in the present case, the statistics can be regarded as those based on a large sample size because the population size in each river was sufficiently large (at least hundreds or thousands individuals; cf. Sakai, 2010, 2011) and eDNA could have captured haplotypes from the whole population. Also, a simulation analysis by scaling down the sample size of each to 2000, 1000 and 500 confirmed that the *F*_ST_ values were still significant in most cases even for the smallest *F*_ST_ between spring and autumn migrants (7-CN; Table S6).

### The spatial genetic structure between the migrant groups

The *F*_ST_ values between spring and autumn migrants in each river were negatively related to latitude and the presence of fish stocking, suggesting that spatial and anthropogenic factors influence the magnitude of genetic differences between the migrant groups. Of these, the latitude of rivers may influence the degree of overlap in the spawning timing between the two migrant groups. We calculated the probability of Ayu spawning at each river every two weeks from the fourth week of August to the second week of November based on the results of 19-year surveys of Ayu spawning in Lake Biwa reported by the Shiga Prefectural Fisheries Experiment Station (See appendix for details, Table S7). As a result, we found that Ayu starts to spawn earlier and continues until later in the spawning season in rivers located at higher latitudes (beta regression model, both the fourth week of August and the second week of November, *p* < 0.05). It is also been hypothesised that autumn migrants begin spawning earlier than spring migrants and that some of them can spawn again roughly 20 days following the first spawning (Shimadu, 1950). As a result, autumn migrants in high-latitude rivers with longer spawning periods are more likely to spawn multiple times. Such multiple spawning by the autumn migrants in higher latitude rivers will increase the chances of simultaneous spawning with spring migrants later in the spawning season, posssibly reflecting the decreased *F*_ST_ value between migrant groups when assuming some homing migration (but see the next section).

The lower *F*_ST_ value between the migrant groups in the rivers where fish stocking has been conducted, on the other hand, could be explained by the admixture of autumn migrant fish into the spring migrant group by the artificial release of the latter in the spring season. In five rivers (9-AD, 8-IS, 4-SR, 3-IN, 1-EC), the juveniles of Ayu caught in Lake Biwa (i.e. not spring migrants) have been stocked into the middle and/or upper reaches every year (Iguchi et al., 2002), leading cohabitation of the two groups during the spring and early summer. Although the proportion and survival until the spawning season of the stocked fish are unclear, the stocked fish may simultaneously spawn with the spring migrants. The overlap in spawning timing and location in this situation could reflect the lower *F*_ST_ between the migrant groups when evaluated by eDNA or even by the conventional capture-based method.

### The spatial and temporal genetic structure within each migrant group

When all rivers were considered, no isolation-by-distance patterns (*F*_ST_ vs. three types of geographic distance) were observed in any migration group. Whereas, when Lake Biwa was divided into east and west sides, a significant association between *F*_ST_ and geographical distance between rivers was observed in the spring migrant samples in 2018 from the west side rivers. Additionally, although statistically not significant probably due to a small sample size (4 rivers) by missing data, a higher Mantel statistic *r* values (*r* > 0.5) were observed in four data sets including both migrant groups. Iguchi et al. (2002) also reported a significant association between *F*_ST_ and the geographical distance based on samples from five rivers located in the west side of the lake [U River (135.99°E, 35.26°N), 9-AD, 7-CN, Oura River (136.12°E, 35.48°N), and 6-SO]. These findings point to the prevalence of weak distance isolation in both migration groups. On the other hand, isolation by distance was not observed when using the shortest straight lines distance between pairs of all the rivers, suggesting that Ayu tends to migrate along the lakeshore instead of taking the route across the center of the lake when the fish search the rivers to enter.

The degree of genetic differentiation among river samples was clearly greater in spring migrants than in autumn migrants. This difference in the genetic structure between the two migrant groups may be explained by the timings of spawning and hatching and by seasonal changes in gyres (i.e. circulating flow of lake water) in Lake Biwa. As previously stated, the landlocked Ayu in Lake Biwa spawns from the end of August to the beginning of November in the lower sections of inlet rivers, and autumn migrants tend to spawn earlier than spring migrants (Azuma, 1973b; Tsukamoto et al., 1987). The eggs hatch in about two to three weeks, and the larval fish immediately flow down to the lake (Azuma, 1970). In Lake Biwa, gyres occur from early summer to autumn, and the largest gyre, called the first gyre, reaches a maximum flow velocity of around 20 cm/s in August and September (Fig. 2, Akitomo et al., 2004). Therefore, the larvae produced by autumn migrants which flow down to the lake earlier (early to middle September) will likely spread more widely across Lake Biwa by still strong gyres. On the other hand, the larvae which were spawned later mainly by spring migrants with some autumn migrants may tend to stay at the river mouth or littoral areas near their home rivers because of the weakend influences of the gyre in autumn. We here propose a hypothesis that the difference in genetic structure between the migrant groups is caused by the differences in the extent of diffusion of the larval fish under seasonal fluctuation of gyres.

Alternative explanation for the difference in genetic structure between the migrant groups would be a homing migration habit presumably enhanced in the spring migrant group. However, this explanation was not supported by the comparisons between two-year samples. The PCoA did not show any tendency for different-year samples from the same rivers to be plotted close together. This result indicates that the genetic structure changed between the two years and does not spport simple homing migration. There are two non-exclusive possiblities to explain this result. One is that the spring migrants are shuffled among rivers every year. The tendency of isolation by distance detected in this study suggests that such shuffling seems to be limited only between neighbouring rivers. However, because Ayu has been inferred to migrate into a nearby river with relatively higher water levels at the time of their migration (Takahashi and Azuma, 2016), fish born in different rivers can be mixed with each other. Another explanation is based on the “swiching hypothesis” proposed by Tsukamoto et al. (1987). This hypothesis was built based on a growth analysis using otoliths and the spawning timing of each migrant group, and involves the change of the migration type between generations. That is, early-born individuals (expected as the progeny of autumn migrants) migrate to the river in the next spring, whereas late-born individuals (expected as the progeny of spring migrants) migrate to the river in the next autumn. Since the autumn migrant group is always far more numerous than the spring migrant group, complete switching of the migration pattern between generations is unlikely. However, even if only the earliest spawners of autumn migrants and the latest spawners of spring migrants switch with each other, the genetic structure would change between 2018 and 2019. To test these hypotheses, examination over several generations is necessary.

## Conclusion

Our results suggested that there were genetic differences between two migrant groups of the landlocked Ayu population inhabiting Lake Biwa, and the *F*_ST_ values were negatively related to latitude and fish release. Additionally, there was the spatial and temporal genetic structure between and within the migrant groups. Although further explicit tests are necessary, genetic differentiation between the migrant groups and geographical population structure were hypothesised to be influenced by differences in spawning timing between populations and lake currents. Further studies on ecological and genetic factors determining the migration timing would give better understanding towards the evolution and maintenance mechanisms of the life history divergence in the Ayu population of Lake Biwa. Finally, as this study has demonstrated, the eDNA approach will be effective for conducting large-scale genetic structural investigations and then contribute as a springboard for further detailed population genetic studies.

## Acknowledgements

We thank Dr. K. Iguchi and Dr. H. Takeshima for providing valuable advice on the work. We also thank Dr. H. Yamanaka, Dr. A. Maruyama and Dr. M. Kondoh for supporting this work as host researchers for the JSPS Research Fellowship for Young Scientists (DC2, S.T.). We thank the fisheries management division in Shiga Prefecture and local fishery associations (Kitafunaki, Momose and Amano) for research permission and supports. This work was supported by Grant-in-Aid for JSPS Research Fellow Grant Number 18J10088 and 21J01519.

## Authors’ contributions

S.T. conceived and designed research. S.T., N.S., and H.S. performed field surveys and molecular analysis. S.T. and K.W performed data analysis. S.T. and K.W. wrote the early draft and it were completed with significant input from all authors.

## Data availability

Full details of the results for each experiment of the present study are available in the Appendix and supporting information (Tables S1–S7). All raw sequences obtained in the qMiFish analysis were deposited in the DDBJ Sequence Read Archive (accession number: DRA013835).

