## Appendix for "Differences in the genetic structure between and within two landlocked Ayu groups with different migration patterns in Lake Biwa revealed by environmental DNA analysis"

[1] Graduate School of Science, Kyoto University, Kitashirakawa-Oiwakecho, Sakyo-ku, Kyoto 606–8502, Japan: [2] Faculty of Science and Technology/Graduate School of Science and Technology, Ryukoku University, 1–5 Yokotani, Seta Oe-cho, Otsu, Shiga 520–2194, Japan: [3] Environmental Research and Solutions co., ltd, Hikaridai 2–3–9, Seika-cho Souraku-gun, Kyoto 619–0237, Japan: [4] Nishinihon Institute of Technology, 9–30 Wakamatsu, Kochi 781–0812, Japan

### Appendix: Extended methodological details

#### *DNA extraction from filter samples for eDNA-based survey*

DNA was extracted from each filter using the spin column (EconoSpin, EP-31201; GeneDesign, Inc., Osaka, Japan) and DNeasy Blood and Tissue Kit (Qiagen, Hilden, Germany) following the procedures described in Tsuji et al. (2020a). The silica-gel membrane which was originally equipped with spin column was removed prior to use. The each GF/F filter was rolled into a cylindrical shape and put into the upper part of the spin column. Spin columns were centrifuged for 1 min at 6,000 g to remove any excess water remaining in the filters (Tsuji et al. 2022). A mixture containing 100 µL of Buffer AL, 10 µL of proteinase K and 200 µL of ultrapure water was added and incubated for 30 minutes at 56ºC. After incubation, the spin columns were centrifuged for 1 min at 6,000 g. To recover any DNA remaining on the filters, 200 µL of Buffer TE (pH 8.0) was added to each filter and incubated for 1 min at room temperature. After centrifuging for 1 min at 6,000 g, the upper part of the spin column was removed, and a mixture containing 100 µL buffer AL and 600 µl ethanol was added to elution and mixed well by pipetting. The mixture was added into a DNeasy Mini spin column and centrifuged for 1 min at 6,000 g. The upper part of the DNeasy Mini spin column was placed on a new 2 mL collection tube, and 500 µL of Buffer AW1 was added. After centrifuging for 1 min at 6,000 g, the upper part of the DNeasy Mini spin column was placed on a new 2 mL collection tube, and 500 µL of Buffer AW2 was added. After centrifuging for 2 min at 15,000 g, the upper part of the spin column was placed on a new 1.5 mL Lobind microcentrifuge tube (Eppendorf, Hamburg, Germany). At the final extraction step, DNA was eluted from the column with 100 µL of Buffer AE. The extracted DNA was stored at −20 ºC until use.

***Paired-end library preparation and sequencing by MiSeq***

To quantitatively evaluate the concentration of each Ayu haplotype, we adopted a quantitative eDNA metabarcoding method using the internal standard DNAs (qMiSeq method; Ushio et al. 2018). In this study, three artificial chimaeric sequences which were designed in Tsuji et al. (2021c) were used as the internal standard DNAs (Table S2). The use of the qMiSeq method enables us to estimate eDNA copy numbers of detected unknown Ayu haplotypes by using sample-specific standard lines which were obtained for each sample based on the relationship between the number of sequence reads and the known copy number of the standard DNAs (Ushio et al. 2018, Tsuji et al. 2021c). In the DNA library preparation, we employed a two-step tailed PCR approach to construct the paired-end libraries. In the first-round PCR, the standard DNA mix that contains the three types of standard DNAs with different copy numbers was added to each reaction solution of PCR: Std. DNA 1 (10 copies/μL), Std. DNA 2 (50 copies/μL) and Std. DNA 3 (100 copies/μL). In the first-round PCR (1st PCR), a species-specific primer set, PaaDlp-2_F and PaaDlp-2_R1/R2 (Tsuji et al. 2020a), was used to amplify the D-loop region of mitochondrial DNA of ayu and the standard DNAs (insert length 166 bp).

1st PCR was carried out with a 12-μl reaction volume containing 6.0 μL of 2 × KAPA HiFi HotStart ReadyMix (KAPA Biosystems, Wilmington, WA, USA), 0.35 μL PaaDlp-2_F (10 μM), 0.175 μL PaaDlp-2_R1/R2 (10 μM), 1.3 μL of sterilised distilled $H_2O$, 1.0 μL of standard DNA mix and 3.0 μL eDNA sample. 1st PCR was performed in five replicates for each eDNA sample, and each PCR plate included five replicates of PCR negative controls to monitor cross-contamination. The thermal cycle profile was as follows: 3 min at 95ºC (initial denaturation) and 35 cycles of 20 s at 98ºC (denaturation), 15 s at 60ºC (annealing), 15 s at 72ºC (extension) and 5 min at 72ºC (final extension). The five replicates of 1st PCR products were pooled for each sample to mitigate the PCR dropouts and purified using Agencourt AMPure XP beads (Beckman Coulter, California, USA) (target amplicon length; ca. 290 bp).

The second-round PCR (2nd PCR) was carried out with a 12-μl reaction volume containing 6.0 μL 2×KAPA HiFi HotStart ReadyMix, 1.4 μL each primer (2.5 μM), 0.2 μL sterile distilled H2O and 3.0 μL the purified 1st PCR product. The primers included the Illumina sequencing adaptors plus the 8-bp identifier indices, and different combinations of indices were used for different templates for a massively parallel sequencing with MiSeq. The thermal cycle profile was as follows: 3 min at 95ºC (initial denaturation) and 12 cycles of 20 s at 98ºC (denaturation), 15 s at 72ºC (combined annealing and extension; shuttle PCR), 5 min at 72ºC (final extension).

All indexed products of the second PCR were pooled in equal volumes. The pooled libraries (total 100 μL) were subjected to agarose gel electrophoresis using 2% E-Gel SizeSelect Agarose Gels (Thermo Fisher Scientific, USA) and excised the target size of the libraries (ca. 370-bp). The DNA concentrations in the excised libraries were estimated using a Qubit dsDNA HS assay kit and a Qubit fluorometer (Life Technologies) and then finally adjusted to 4 nM (assuming 1 bp of DNA has the molecular weight of 660 g $mol^{-1}$). The adjusted library was sequenced on the MiSeq platform using a MiSeq v2 Reagent Micro or Nano Kit for 2 × 150 bp PE cartridge (Illumina, San Diego, USA) with 30% PhiX spike-in according to the manufacturer's instructions.

***Probability of Ayu spawning based on the result of visual surveys by the Shiga Prefectural Fisheries Experiment Station***

We calculated the probability of Ayu spawning at each river every two weeks from the fourth week of August to the second week of November based on the results of 19-year visual surveys of Ayu spawning in Lake Biwa reported by the Shiga Prefectural Fisheries Experiment Station. A total of 11 rivers were surveyed, including nine rivers (1-EC, 3-IN, 4-SR, 5-AM, 6-SO, 7-CN, 8-IS, 9-AD and 11-WD) that were also surveyed in this study and the other two rivers (Yasu River; 35°05'50.0"N, 135°59'36.1"E and Ane River; 35°23'33.8"N, 136°13'21.2"E). The probability of spawning at each survey timing was calculated as the average over 19 years when the presence or absence of eggs was expressed as 1 or 0 (Table S7). A beta regression model was used to investigate the relationships between the latitude of each river mouth and the probability of spawning at early (fourth week of August) or end (second week of November) of the spawning season (Fig. S1).


### Figure

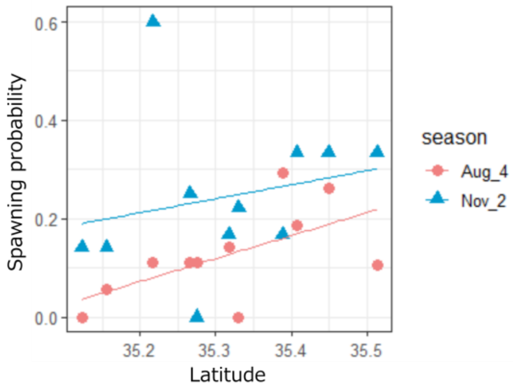


Fig. S1 The relationships between the latitude of each river mouth and the probability of spawning at the early (fourth week of August) or end (second week of November) of the spawning season. Solid lines indicate linear regression lines (all lines are significant; beta regression model, $p < 0.05$).
